# Final amendment: A plausible explanation for *in silico* reporting of erroneous MET gene expression in tumor-educated platelets (TEP) intended for "liquid biopsy" of non-small cell lung carcinoma still refutes the TEP-study

**DOI:** 10.1101/148718

**Authors:** Sandeep Chakraborty

**Affiliations:** R - 44/ 1, Celia Engineers, T. T. C Industrial Area, Rabale, Navi Mumbai, 400701, India

## Abstract

**Final amendment note:** This paper had proposed a plausible way for detecting large quantities of MET, which the authors have clarified was not done :the possible explanation proposed for this erroneous MET gene expression does bypass the filtering step we perform in the data processing pipeline, i.e. selection of intron-spanning reads, as can be read in the main text” comments in http://www.biorxiv.org/content/early/2017/07/02/146134, where a continuing critique of the TEP study continues. Please consider this pre-print closed.

**Original abstract:** The reported over-expression of MET genes in non-small cell lung carcinoma (NSCLC) from an analysis of the RNA-seq data from tumor-educated platelets (TEP), intended to supplement existing ‘liquid biopsy’ techniques [1], has been refuted recently (http://biorxiv.org/content/early/2017/06/05/146134, not peer-reviewed). The MET proto-oncogene (Accid:NG 008996.1, RefSeqGene LRG 662 on chromosome 7, METwithintrons) encodes 21 exons resulting in a 6710 bps MET gene (Accid: NM 001127500.2, METonlyexons). METwithintrons has multiple matches in the RNA-seq derived reads of lung cancer samples (for example: SRR1982756.11853382). Unfortunately, these are non-specific sequences in the intronic regions, matching to multiple genes on different chromosomes with 100% identity (KIF6 on chr6, COL6A6 on chr3, MYO16 on chr13, etc. for SRR1982756.11853382). In contrast, METonlyexons has few matches in the reads, if at all [2]. However, even RNA-seq from healthy donors have similar matches for METwithintrons so the computation behind the over-expression statistic remains obscure, even if METwithintrons was used as the search gene. In summary, this work re-iterates the lack of reproducibility in the bioinformatic analysis that establishes TEP as a possible source for “liquid biopsy”.

## Introduction

Tumor tissue biopsy, the gold standard for cancer diagnostics, pose challenges that include access to the tumor, quantity and quality of tumoral material, lack of patient compliance, repeatability, and bias of sampling a specfic area of a single tumor [3]. This has resulted in a new medical and scientific paradigm defined by minimal invasiveness, high-efficiency, low-cost diagnostics [4], and, whenever possible, personalized treatment based on genetic and epigenetic composition [5]. The presence of fragmented DNA in the cell-free component of whole blood (cfDNA) [6], first reported in 1948 by Mandel and Metais, has been extensively researched for decades, with extremely promising results in certain niches [7]. Additionally, cfDNA derived from tumors (ctDNA) [8] have tremendous significance as a cancer diagnostic tool [9], and for monitoring responses to treatment [10]. However, detection of ctDNA, and differentiation with cfDNA, remains a challenge due the low amounts of ctDNA compared to cfDNA [11].

Recently, tumor-educated blood platelets (TEP) were proposed as an alternative source of tumor-related biological information [1, 12]. The hypothesis driving the potential diagnostic role of TEPs is based on the interaction between blood platelets and tumor cells, subsequently altering the RNA profile of platelets [13, 14]. The study showed using RNA-seq data that tumor-educated platelets (TEP) can distinguish 228 patients with localized and metastasized tumors from 55 healthy individuals with 96% accuracy [1]. As validation, this study reported significant over-expression of MET genes in non-small cell lung carcinoma (NSCLC), and HER2/ERBB2 [15] genes in breast cancer, which are well-established biomarkers.

Previously, the TEP-study was refuted by an analysis of a subset of the samples (yet to be peerreviewed) [2]. Here, an analysis based on the complete MET gene (both introns and exons, Accid:NG 008996.1) demonstrates that intronic non-specific sequences might mislead bioinformatic analysis. Moreover, considering that RNA-seq from healthy donors have similar matches with the complete MET gene, the computation behind the over-expression statistic remains obscure [1].

## Results

The MET proto-oncogene (Accid:NG 008996.1, RefSeqGene LRG 662 on chromosome 7, METWITHINTRONS) encodes 21 exons leading to a 6710 bps MET gene (Accid: NM 001127500.2, METONLYEXONS). METWITHINTRONS has multiple matches in the RNA-seq derived reads (for example: SRR1982756.11853382) (Fig. 1). Unfortunately, these are non-specific sequences in the intronic regions: SRR1982756.M.11853382(CTTCACGTAGTTCTCGAGCCTTGGTTTTCAGCTCCATCAGCTCCTTTAAGCACTTCTCTGTA TTGGTTATTCTAGTTATACATTCTTCTAAATTTTTTTCA) matches to mutliple genes on different chro-mosomes (KIF6 on chr6, COL6A6 on chr3, MYO16 on chr13, etc). In contrast, METONLYEXONS has few matches in the reads, if at all [2]. Thus, it is erroneous to assign the intronic sequences being expressed to the MET gene. However, even RNA-seq from healthy donors have similar matches for METWITHINTRONS (Fig 3) - so the computation behind the over-expression statistic remains obscure, even if METwithintrons was used as the search gene.

**Figure 1:**
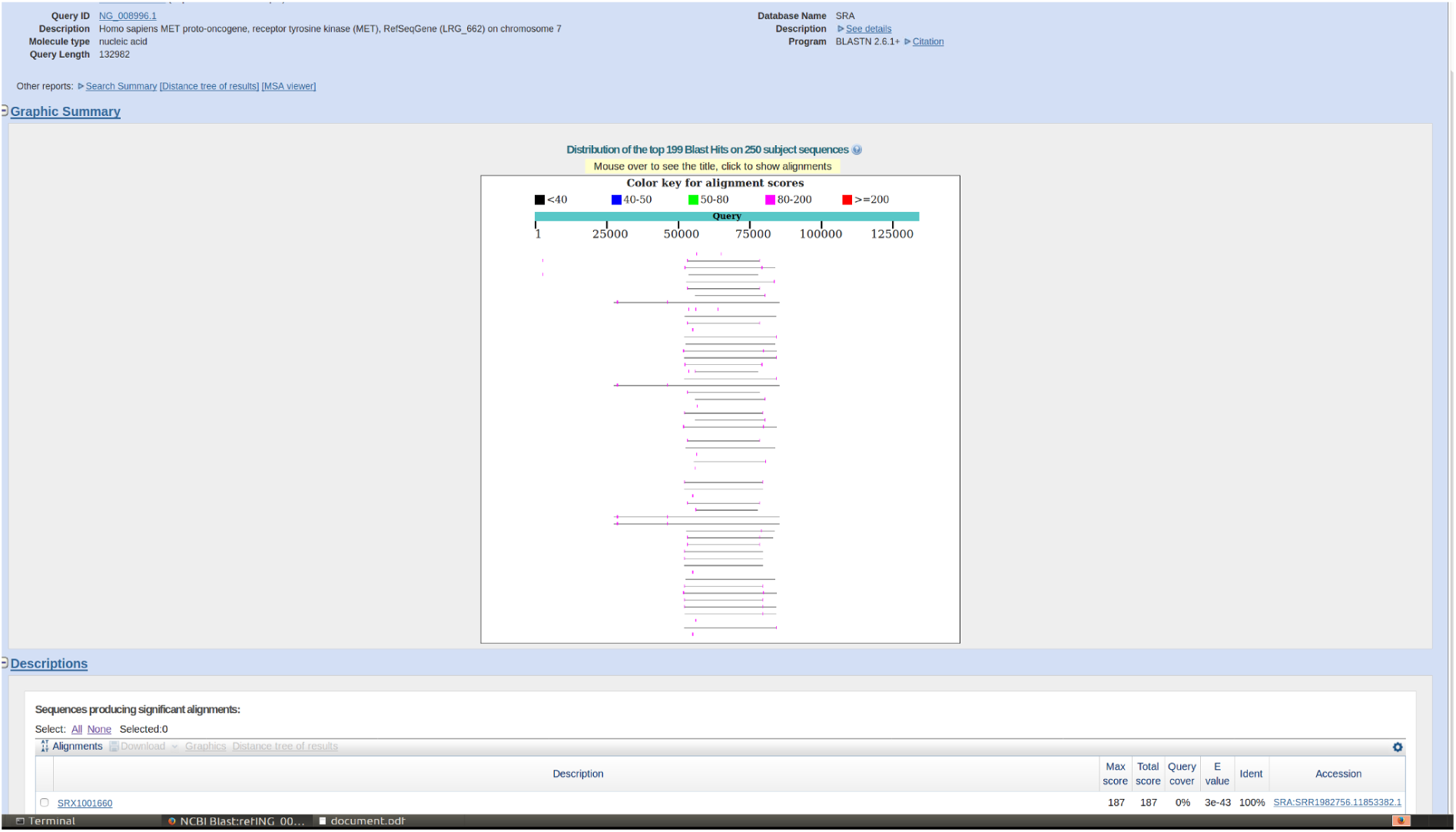
Graphical representation of matches to the complete MET gene (including introns) to a lung cancer sample (SRR1982756): This shows the first 100 matches, out of thousands of significant matches using the online BLAST interface. However, as demonstrated previously [2], very few of these matches are in the exonic region. Also, these sequences (an example is SRR1982756.11853382) are very non-specific and match to many genes in different chromosomes (Fig 2).

**Figure 2:**
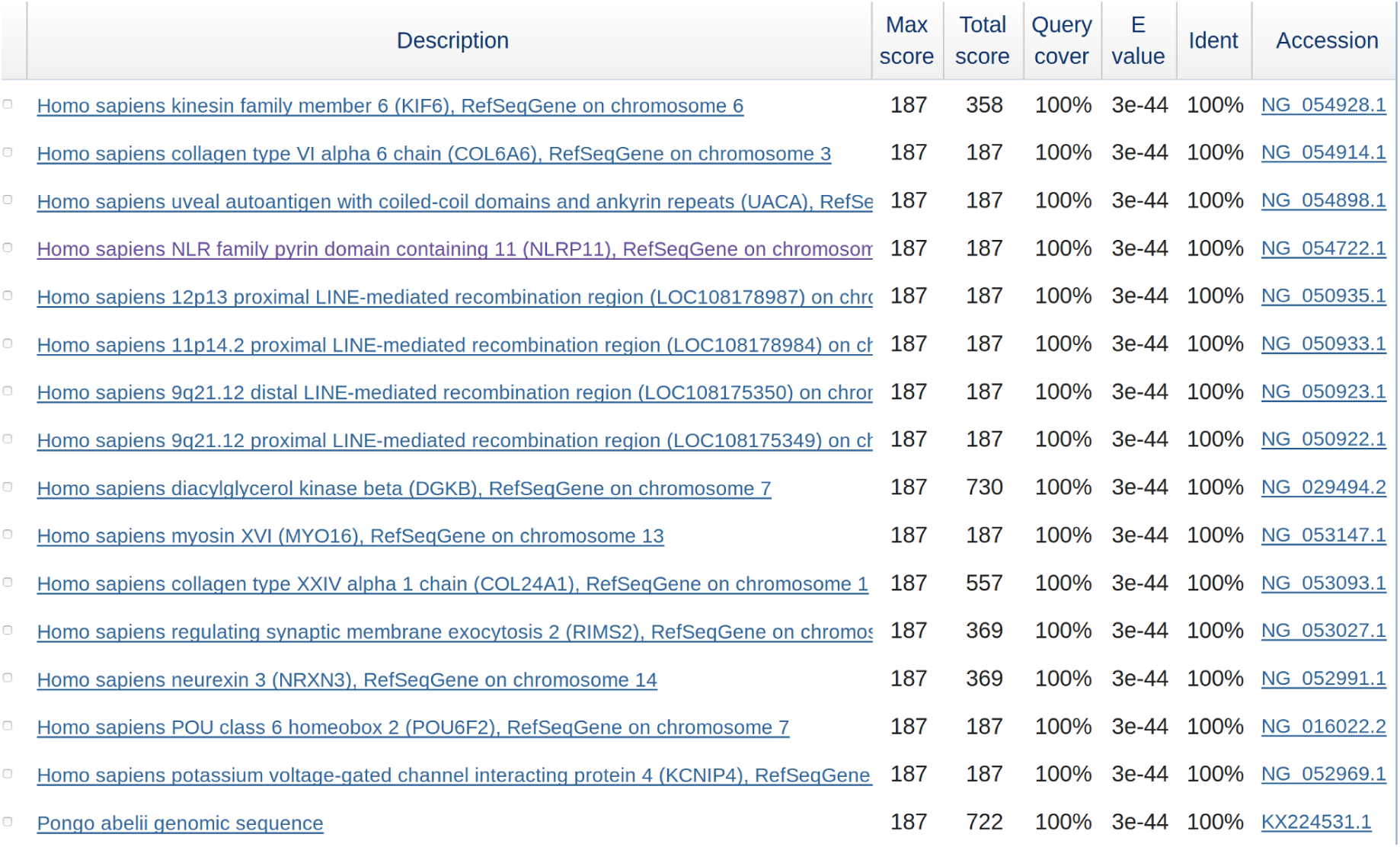
Non-specific nature of the intronic sequences of the MET gene that match to the RNA-seq reads from the platelets: The sequence SRR1982756.M.11853382 matches to several genes across different chromosomes - KIF6 on chr6, COL6A6 on chr3, MYO16 on chr13, etc.

**Figure 3:**
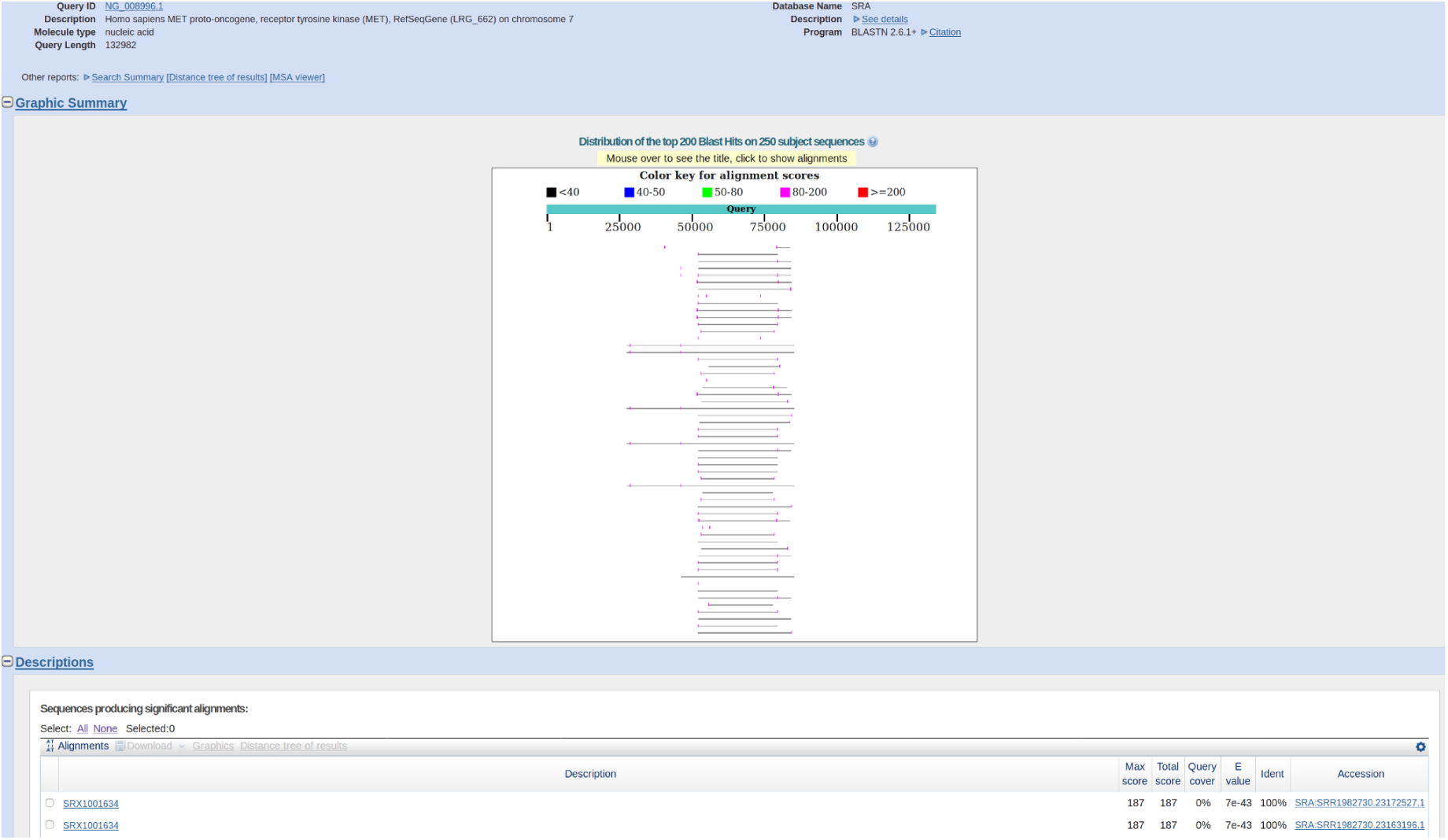
Graphical representation of matches to the complete MET gene (including introns) to a healthy donor sample (SRR1982730): This shows the first 100 matches, out of thousands of significant matches, as found in lung cancer samples as well (Fig. 1). Thus, any statistic showing over-expression must be validated against these raw numbers. These sequences are very non-specific and match to many genes in different chromosomes (Fig 2).

## Conclusion

Here, the absence of MET over-expression as reported in the TEP-study [1] is investigated in further detail by using the full MET gene including introns. It turns out that several intronic sequences have matches in the RNA-seq samples. However, these intronic sequences are non-specific (i.e. matching to several other genes with 100% identity). Further, there is to be large number of matches in healthy donor samples as well. This work re-iterates the lack of reproducibility in the bioinformatic analysis that establishes TEP as a possible source for “liquid biopsy”.

## Materials and methods

The BLAST interface suffices to demonstrate the presence of non-specific sequences from the intronic regions of the MET gene to the RNA-seq samples, and the non-specific nature of these sequences based on 100% identity to a plethora of genes in different chromosomes. These have been verified by a kmer-based version (KEATS [16]) of YeATS [17–21], as well.

**Table 1:** Raw counts of reads matching to the MET gene: Although this is a subset (24 out of 60 NSCLC, and 20 out of 60 healthy), and the numbers are not normalized, its seems unlikely that any statistic will show MET over-expression in NSCLC. It does not even add up to one complete gene in most cases.

|  |  |  |  |
| --- | --- | --- | --- |
| Healthy | SRR1982752 | NM_001127500.2 | ntranscripts=278+2 |
|  | SRR1982731 | NM_001127500.2 | ntranscripts=194+0 |
|  | SRR1982742 | NM_001127500.2 | ntranscripts=56+2 |
|  | SRR1982702 | NM_001127500.2 | ntranscripts=127+2 |
|  | SRR1982750 | NM_001127500.2 | ntranscripts=174+2 |
|  | SRR1982741 | NM_001127500.2 | ntranscripts=124+2 |
|  | SRR1982720 | NM_001127500.2 | ntranscripts=7+0 |
|  | SRR1982730 | NM_001127500.2 | ntranscripts=171+0 |
|  | SRR1982722 | NM_001127500.2 | ntranscripts=2+0 |
|  | SRR1982735 | NM_001127500.2 | ntranscripts=132+0 |
|  | SRR1982740 | NM_001127500.2 | ntranscripts=179+2 |
|  | SRR1982751 | NM_001127500.2 | ntranscripts=152+2 |
|  | SRR2095004 | NM_001127500.2 | ntranscripts=23+0 |
|  | SRR1982721 | NM_001127500.2 | ntranscripts=2+0 |
|  | SRR1982732 | NM_001127500.2 | ntranscripts=180+2 |
|  | SRR1982737 | NM_001127500.2 | ntranscripts=167+2 |
|  | SRR1982700 | NM_001127500.2 | ntranscripts=78+2 |
|  | SRR2095014 | NM_001127500.2 | ntranscripts=23+0 |
|  | SRR1982710 | NM_001127500.2 | ntranscripts=28+0 |
|  | SRR1982738 | NM_001127500.2 | ntranscripts=69+2 |
|  | SRR1982701 | NM_001127500.2 | ntranscripts=99+3 |
| NSCLC | SRR1982781 |  |  |
|  | SRR1982780 | NM_001127500.2 | ntranscripts=13+2 |
|  | SRR1982791 | NM_001127500.2 | ntranscripts=24+0 |
|  | SRR2096517 | NM_001127500.2 | ntranscripts=5+0 |
|  | SRR1982772 | NM_001127500.2 | ntranscripts=8+4 |
|  | SRR1982770 | NM_001127500.2 | ntranscripts=8+2 |
|  | SRR1982790 | NM_001127500.2 | ntranscripts=44+2 |
|  | SRR2096502 | NM_001127500.2 | ntranscripts=1+0 |
|  | SRR1982756 | NM_001127500.2 | ntranscripts=5+0 |
|  | SRR1982759 | NM_001127500.2 | ntranscripts=10+0 |
|  | SRR1982795 | NM_001127500.2 | ntranscripts=9+0 |
|  | SRR1982762 | NM_001127500.2 | ntranscripts=11+2 |
|  | SRR1982761 | NM_001127500.2 | ntranscripts=5+0 |
|  | SRR2096503 | NM_001127500.2 | ntranscripts=2+0 |
|  | SRR2096501 | NM_001127500.2 | ntranscripts=33+2 |
|  | SRR1982782 | NM_001127500.2 | ntranscripts=2+2 |
|  | SRR2096516 | NM_001127500.2 | ntranscripts=6+2 |
|  | SRR1982777 | NM_001127500.2 | ntranscripts=40+2 |
|  | SRR1982793 | NM_001127500.2 | ntranscripts=6+0 |
|  | SRR1982765 | NM_001127500.2 | ntranscripts=16+0 |
|  | SRR1982771 | NM_001127500.2 | ntranscripts=11+2 |
|  | SRR1982787 | NM_001127500.2 | ntranscripts=2+0 |
|  | SRR1982792 | NM_001127500.2 | ntranscripts=5+0 |
|  | SRR1982760 | NM_001127500.2 | ntranscripts=2+0 |

## Competing interests

No competing interests were disclosed.

